# Song sparrows fail to discriminate between current and historical songs

**DOI:** 10.64898/2026.07.02.736160

**Authors:** William A. Searcy, Susan Peters, Gabriel Macedo, Stephen Nowicki

## Abstract

Song evolves rapidly in songbirds, as has been proven for a substantial number of songbird species by demonstrating acoustic differences between current songs and "historical" songs recorded 20 or 30 years previously. In two species of songbirds, white-crowned sparrows and savannah sparrows, it has been further shown that evolutionary changes over such time spans are sufficient to affect the response of receivers, with territorial males responding more aggressively to current songs than to historical ones. These two species, however, have especially low population variability in song, with most males in any population singing the same, single song type; this background of song uniformity makes it especially easy to discern temporal changes. Here we examine response to temporal change in song in a third sparrow species, the song sparrow, in which population variation in song is much higher: males sing 5-13 song types each, with low song-sharing between males, so that hundreds of distinct song types occur in a local population. We find evidence that temporal change has occurred in the songs of our study population, in that 24 song current song types share more introductory phrases with other current song types than do 24 historical song types recorded 27-29 years earlier. Nevertheless, in a song playback experiment, current males showed no difference in response to current and historical songs. The results are in accord with the hypothesis that high levels of population variability in song make temporal changes in song difficult to discern.

**Significance statement:** The song of songbirds is shaped by social learning, making birdsong a model system for the study of cultural evolution. Because cultural evolution is thought to occur more rapidly than genetic evolution, many studies have examined changes in song over time periods thought too short to allow measurable genetic evolution. Here we show that song has evolved in our study population of song sparrows, with the relative abundance of introductory phrases changing over a period of just 27-29 years. We find, however, that the song sparrows themselves fail to discern these changes, responding just as strongly to the old ("historical") song types as to current ones. Measurable cultural evolution thus occurs but without functional consequences.

## Introduction

The development of songbird song depends on social learning (Marler 1970; Beecher and Brenowitz 2005; Catchpole and Slater 2008), and birdsong has therefore become a model in the study of cultural evolution (Whiten 2019). Cultural evolution is assumed to occur more rapidly than genetic evolution (Richerson et al. 2010; Perreault 2012), an assumption that has encouraged studies examining evolution of song over periods of time usually thought too short to allow for appreciable genetic evolution. Many of these studies have found evidence of changes in song at the population level over periods of 20 or 30 years or even less. In some cases, the changes observed appear to be rather small (Harbison et al. 1999; O’Loghlen et al. 2011; Graham et al. 2021), but in others major changes have been found in the relative abundance of different song types (Payne 1985; Byers et al. 2010; Ju et al. 2019; Souriau et al. 2025) or in average acoustic measures of song (Luther and Derryberry 2012; Williams et al. 2013; Otter et al. 2020). Whether the observed changes have functional importance is an additional question, one that a small number of studies have addressed by comparing the behavioral response of contemporary conspecifics to current versus "historical" songs.

Significant differences in response to current and historical songs were first demonstrated in studies of white-crowned sparrows (*Zonotrichia leucophrys*) (Derryberry 2007; Derryberry 2011; Luther and Derryberry 2012) and have also been demonstrated in savannah sparrows (*Passerculus sandwichensis*) (Williams et al. 2022). A test for such discrimination by thrush nightingales (*Luscinia luscinia*) gave negative results (Ivanitskii et al. 2025). Here we test for differential response to current and historical songs in a fourth species, the song sparrow (*Melospiza melodia*), which exhibits a more complex pattern of song variability at the population level than the previously-tested species.

White-crowned sparrows show particularly little variability in song structure at the within-population level: almost all males sing only a single song type, and for the most part all the males within a local population produce the same type (Marler and Tamura 1962; Baptista 1977). Against this background of within-population uniformity, it is fairly easy for researchers to discern differences in song between temporally separate samples from a single geographic area (Baptista and Gaunt 1997; Nelson et al. 2004) or between contemporary populations in different geographic areas (Marler and Tamura 1962; Baptista 1977; Baptista and King 1980; Podos and Warren 2007). White-crowned sparrows themselves have demonstrated temporal discrimination between songs recorded at the same sites 24 years apart (Derryberry 2007), 27 years apart (Derryberry 2011), and 36 years apart (Luther and Derryberry 2012). This species has also demonstrated geographic discrimination (Milligan and Verner 1971; Nelson and Soha 2004) including in cases in which the tested dialects are from contiguous areas whose centers are 12 km apart or less (Baker 1983; Tomback et al. 1983).

Savannah sparrows, a second species in which temporal discrimination of songs has been demonstrated, also show low within-population variability in song. Almost all males in this species sing only a single song type (Borror 1961; Wheelwright et al. 2008), and the males within a locality tend to share one predominant type, though with more between-male variation than seen in white-crowned sparrows (Borror 1961; Pitocchelli 1981). Songs have been shown to vary geographically, both at the scale of hundreds of kilometers or more (Chew 1981; Bradley 1994) and at the scale of hundreds of meters or less (Hensel et al. 2022). In a playback study, males of this species responded more strongly to local song elements than to foreign elements recorded approximately 500 km distant (Williams et al. 2019). Evolutionary changes in savannah sparrow song have been found over periods of approximately 15 years (Bradley 1994), 30 years (Williams et al. 2013), and 27 years (Williams et al. 2022). Williams et al. (2022) showed via a playback experiment that savannah sparrows of both sexes discriminate between artificial songs that mimic changes - in the presence and number of a particular note type - that occurred in the study population over approximately 27 years.

Thrush nightingales, the third species in which temporal discrimination has been tested, have considerably larger song repertoire sizes than white-crowned and savannah sparrows, with average repertoire sizes ranging from 6 to 24 song types per male in various populations (Griessman and Naguib 2002, Ivanitskii et al 2025, Souriau et al 2025). Song type sharing is common between males within locales (Griessman and Naguib 2002; Souriau et al. 2025), but even so the overall diversity of songs within populations is undoubtedly much greater than in the two sparrow species. Geographic variation in song occurs, in that song type sharing falls off with distance between populations (Sorjonen 1987), but we know of no experimental tests for geographic discrimination in this species. Changes in the song population are facilitated in this species by the fact that individuals are capable of changing their repertoires as adults (Sorjonen 1987). Rapid temporal change in song type usage has been documented, with a 90% turnover in song types over 16 years in a Finnish population (Sorjonen 1987) and two complete turnovers in a Russian population, the first in 39 years and the second in five years (Ivanitskii et al. 2025). Relatively rapid change in the acoustic structure of individual song types has also been observed (Ivanitskii et al. 2023). Despite these considerable temporal changes in song, a playback study with males of this species failed to find discrimination of song samples recorded 48 years apart (Ivanitskii et al. 2025).

Greater within-population song diversity has been suggested to make discrimination of geographic variation more difficult (Searcy et al. 2003), and it seems logical that this difficulty would also carry over to discrimination between current and historical songs. The results obtained so far on temporal discrimination support this hypothesis: two species with low within-population song diversity (white-crowned sparrows and savannah sparrows) discriminate between current and historical song, whereas a species with higher within-population song diversity (thrush-nightingales) does not. Here we further test this hypothesis by investigating temporal discrimination in song sparrows. Repertoire sizes in this species average 8-12 song types per male, depending on the population (Hill et al. 1999; Peters et al. 2000; Hughes et al. 2007), considerably larger than in white-crowned and savannah sparrows but smaller than in thrush nightingales. Importantly, song sparrows have very low levels of song sharing in eastern populations (Hughes et al. 1998); Searcy et al. 2014), though song sharing is considerably higher in western populations (Hill et al. 1999). In our eastern study population, males share on average only 8.2% of their song types with each immediate neighbor (DuBois et al. 2016), and sharing is expected to be even lower with non-neighbors (Hill et al. 1999). With song sharing so low, the total number of distinctive song types increases rapidly as greater numbers of neighboring males are examined. Borror (1961), for example, estimated that there were close to 200 distinct song types in the repertoires of 46 song sparrows occupying a 16-hectare plot in Maine. Population-wide song diversity thus seems even greater in song sparrows than in thrush nightingales (Ivanitskii et al. 2023).

Song sparrow songs have been shown to vary geographically, both in the occurrence of specific song elements (Harris and Lemon 1974) and in average acoustic measurements (Patten et al. 2004). A number of studies have shown that song sparrows discriminate geographic variation (Harris and Lemon 1974; Searcy et al. 1997; Searcy et al. 2002; Patten et al. 2004), but, to our knowledge, song sparrows have not previously been tested for temporal discrimination. Here we test male song sparrows at our Pennsylvania study sites for discrimination between local songs recorded in 1992-1994 and songs recorded at the same sites in 2021, a gap of 27-29 years. The experimental design is modeled on a previous study of geographic discrimination in this species (Searcy et al. 1997), in which males at these same Pennsylvania sites discriminated in their territorial responses between local songs and songs from New York sites approximately 540 km distant. We also test whether the song type and phrase type composition of our study population changed over time in a way that might affect response to song.

## Methods

### Study Sites and Subjects

The study was carried within a 6 km radius of a point (41.59 N, 80.35 W) between Linesville and Hartstown in northwestern Pennsylvania, U.S.A. Within this area, we used sites managed by the Pennsylvania Game Commission, Pymatuning State Park, and the Pymatuning Laboratory of Ecology. Subjects were male song sparrows holding territories in old fields or along the margins of lakes and ponds within this area.

### Song stimuli and playback

We used territorial playback (Weeden and Falls 1959) to test male song sparrows for discrimination between current and historical songs. In designing this experiment, we replicated the exact methods used by Searcy et al. (1997) to test for discrimination between Pennsylvania and New York songs.

The historic songs used in the playback experiment were recorded in the study area in 1992-1994. The recordings were made with Marantz PMD 221 and Sony TCM 5000EV cassette tape recorders with either a Sennheiser ME88 shotgun microphone or a Sony ECM-170 microphone in a Sony PBR-330 parabola. These recordings were later digitized at a 44.1 kHz sample rate using Raven Pro (Cornell Laboratory of Ornithology). The current songs were recorded in the same area in 2021 using Marantz PMD 660 and 670 digital recorders and Shure SM58 cardioid microphones in Sony PBR-330 parabolas at a sampling rate of 44.1 or 48 kHz.

We resampled the current songs to match the historical song sample rate using the “resamp” function in Signal v.4 (Engineering Design, Berkeley, CA). We normalized each song to a peak amplitude of 2.0 V and high-pass filtered them at 1500 Hz, also using Signal v.4.

Next, because the historical and current songs were recorded at different times using different equipment, two listeners (SN and Jill Soha) who were blind to treatment (i.e., historical versus current) listened under headphones to all songs presented in a random order, and subjectively rated each song as having low, moderate, or high background noise. This analysis revealed a perceptible difference in noise levels, with historical songs being ranked as slightly noisier than current songs (p = 0.004, two sample t-test assuming equal variances). To adjust for this difference, we selected 4 secs of background sound within an interval of about 10 secs occurring before or after each historical and current song. We then used linear addition of the sound files in Signal v.4 to add historical background sound to its current song playback counterpart (historical and current playback tracks were paired, see below), and vice versa.

Thus, all songs received background sound associated with the songs presented in the counterpart of the stimulus pair. Following this adjustment, the same two listeners again rated the background noise level of each song, still blind to treatment and with songs presented randomly, and no significant difference was detected.

Twelve pairs of current and historical stimulus sets were used for playback, with each pair played to two males, giving a total sample size of 24 subjects. For each subject, we randomly chose either the current or the historic stimulus to present first and presented the other stimulus two or more days later. Each stimulus set included two song types from the appropriate era. For playback, the songs were spaced at intervals of one per 10 sec, in the order: 9 repetitions of type A, 9 type B, 9 type A, and 9 type B. The total playback time thus was 6 min.

Playback trials were carried out from May 11 to May 29, 2025 between 6:00 and 10:00 AM. Playback stimuli were played from a Marantz PMD 660 recorder over an Anchor Audio AN MINI speaker, at an average amplitude of 86 dB SPL (range 82–89; RadioShack 33-2055 digital sound level meter, A-weighting, slow response). This amplitude range largely overlaps with calibrated microphone measurements made in the field of singing song sparrows in this population (Anderson et al. 2008). The speaker was placed face-up near the center of a male’s territory. The position was marked with flagging, and the same position was used on the second test with that male. During playback, the sole response measure noted was the subject’s distance to the speaker, which has been shown to be a reliable predictor of physical attack in song sparrows (Searcy et al. 2006) and which was the sole measure taken in the geographic discrimination study (Searcy et al. 1997). Two observers (WAS and SN) were present in all trials, both helping to keep track of the subject’s position relative to the speaker. In all trials, both observers were blind to which treatment (current or historical) was being presented. To aid in estimating distances, we set out flagging prior to each trial at distances of 2m, 4m, 8m, and 16m on either side of the speaker. Distances were noted in five-second increments in categories of < 2m, 2 to 4m, 4 to 8m, 8 to 16m, and >16m. Observations were started when the first playback song was played, continued through the six-min playback period, and then through a three-min post-playback period. Mean distance was calculated following the method of (Peters et al. 1980) for three-min blocks: the first three minutes of playback, the second three minutes of playback, and a three-min post-playback period.

### Song similarity analysis

As a test of whether measurable changes occur in the songs of the song sparrow population over time, we examined sharing between the two sets of playback songs and the most current recordings we had of full repertoires of our study population males. These full repertoire recordings were from 15 males recorded in 2021 and five males recorded in 2025 (using the same equipment as in 2021). The 2021 recordings were the source of the current playback songs. If the song population as a whole changed between the historic period (1992-1994) and the current period (2021-2025), then song-sharing between the historic playback songs and the current population should be lower than song-sharing between the current playback songs and the current population. Song sharing was judged using spectrograms produced with Audacity v.3.3.3 using a Hann window, a 2048 window size, and a 0-10 kHz frequency axis. The criteria we used for whole-song sharing were those proposed by Hill et al. (1999) for song sparrow songs, which have since been used in other studies (Foote and Barber 2007; Searcy et al. 2014; DuBois et al. 2016). Here, two songs are considered to be shared if they match in two-thirds or more of their component phrases. Phrases are of two main types: trills of repeated syllables and note complexes consisting of unrepeated notes. Trills are considered to match if their syllables are similar in shape, frequency and timing. Note complexes match if half or more of their notes are similar in shape and frequency. In borderline cases, similarity in introductory elements is weighted more strongly than later parts of the song.

We also analyzed sharing of introductory phrases between the two sets of playback songs and current repertoires. Whole song sharing is so low between contemporary individuals in our population (Hughes et al. 1998; DuBois et al. 2016) that it would be difficult for whole song sharing with a historical population to be significantly lower. Sharing of introductory phrases between contemporary individuals in our population is considerably higher (Hughes et al. 1998; DuBois et al. 2016) and is relevant because introductory phrases are especially important to how song sparrows themselves classify song types (Horning et al. 1993; Burt et al. 2002; Anderson et al. 2005). Thus, an analysis of whether the historic playback songs are less likely to share introductions with the current song population than are the current playback songs should be relevant to whether the population of songs has changed over time in a way that might be salient to song sparrows.

## Statistical analysis

For the analysis of song type sharing between the historical playback songs and the current song repertoires, we compared each of the 24 historical playback songs to each of the 154 song types found in the repertoires of the 20 males in the current population sample. For the analysis of song type sharing between the current playback songs and the current song repertoires, we compared each of the 24 current playback songs to the 142-149 song types found in current repertoires other than the repertoire that the focal playback song was taken from. The proportion of current songs matched for whole type or for introduction was then compared between the historical and current playback songs with Mann-Whitney U tests.

Playback data were analyzed in R 4.5.2. We assessed the effect of playback treatment (historical vs. current) and trial order (first vs. second) on the birds’ approach response with Bayesian multilevel models using the R packages *brms* (Bürkner 2017; Bürkner et al. 2024) and *bayestestR* (Makowski et al. 2019). As response variables, we specified, one at a time, the distance from the loudspeaker in three 3-minute periods (first 3-min of playback, second 3-min of playback, 3-min post-playback) as well as the total 9-min (Searcy et al. 1997). We specified individual ID and recording ID as group-level effects to account for repeated measures and thus avoid pseudoreplication at these levels. We specified weakly informative normal priors for the explanatory variables and intercepts (*N* ∼ 0, 1; mean, standard deviation), gamma priors for group-level and residual standard deviations (*gamma* ∼ 1,1; rate, shape), and a lognormal distribution for the response variables, as this distribution fit better than the normal and skew normal distributions. We evaluated model fit and convergence using leave-one-out cross validation, posterior predictive checks, and the scale reduction factor (Gelman and Rubin 1992). We ran all models in four Markov chains, 10,000 iterations (with 50% used in warm-up), and a thinning rate of one (Link and Eaton 2012).

Because our sample size is relatively small (24 individuals, each tested twice), we compared the results of the Bayesian models with those of non-parametric kernel regressions using the R package *np* (Racine and Hayfield 2026) assessing significance (P < 0.05) with 1,000 bootstrap replicates.

## Results

### Song similarity

Whole song sharing between the historical playback songs and the current song repertoires was low (mean proportion of repertoire song types matched = 0.00050), with 22 of 24 historical playback songs matching 0 songs in the current song comparison group of 154 songs. Whole song sharing also was low between the current playback songs and the current repertoires (mean = 0.00171), with 19 of 24 current playback songs matching 0 songs among the 142-149 current songs to which they were compared. A Mann-Whitney U test showed no significant difference in the distribution of whole song sharing proportions for the two playback sets (U = 247, N = 48, P = 0.168).

Sharing of song type introductions was higher and gave evidence of a change in the song population between the historical and current periods. The mean proportion of introductions shared with current song types was 0.00154 for the historical playback songs and 0.00833 for the current playback songs, and the difference in proportion shared values was significant by a Mann Whitney U test (U = 192, N = 48, P = 0.018). Nevertheless, the majority of playback songs in both groups shared introductions with 0 of the current comparison songs (19 of 24 for historical playback songs, 13 of 24 for current playback songs).

### Playback experiment

Responses were similar to playback of historical and current songs for the two three-minute playback periods and for the three-minute post-playback period, as well as for the total 9-minute trial time (Figure 1). This similarity can be seen from the near congruence of the median responses and the wide overlap of the response distributions in each of the three trial periods and in the total trial time (Figure 1). The Bayesian multilevel models give estimates of playback treatment effects that are well within the 95% confidence intervals surrounding 0 for all three trial periods and for the total trial time (Table1), thus indicating no credible effects of playback treatments. The Bayesian models control for trial order, but that factor also appears to have had no credible effects on response (Table 1). The non-parametric analysis agreed with the Bayesian models in indicating no credible effects on response of either playback treatment or trial order (Table 1).

**Figure 1.**
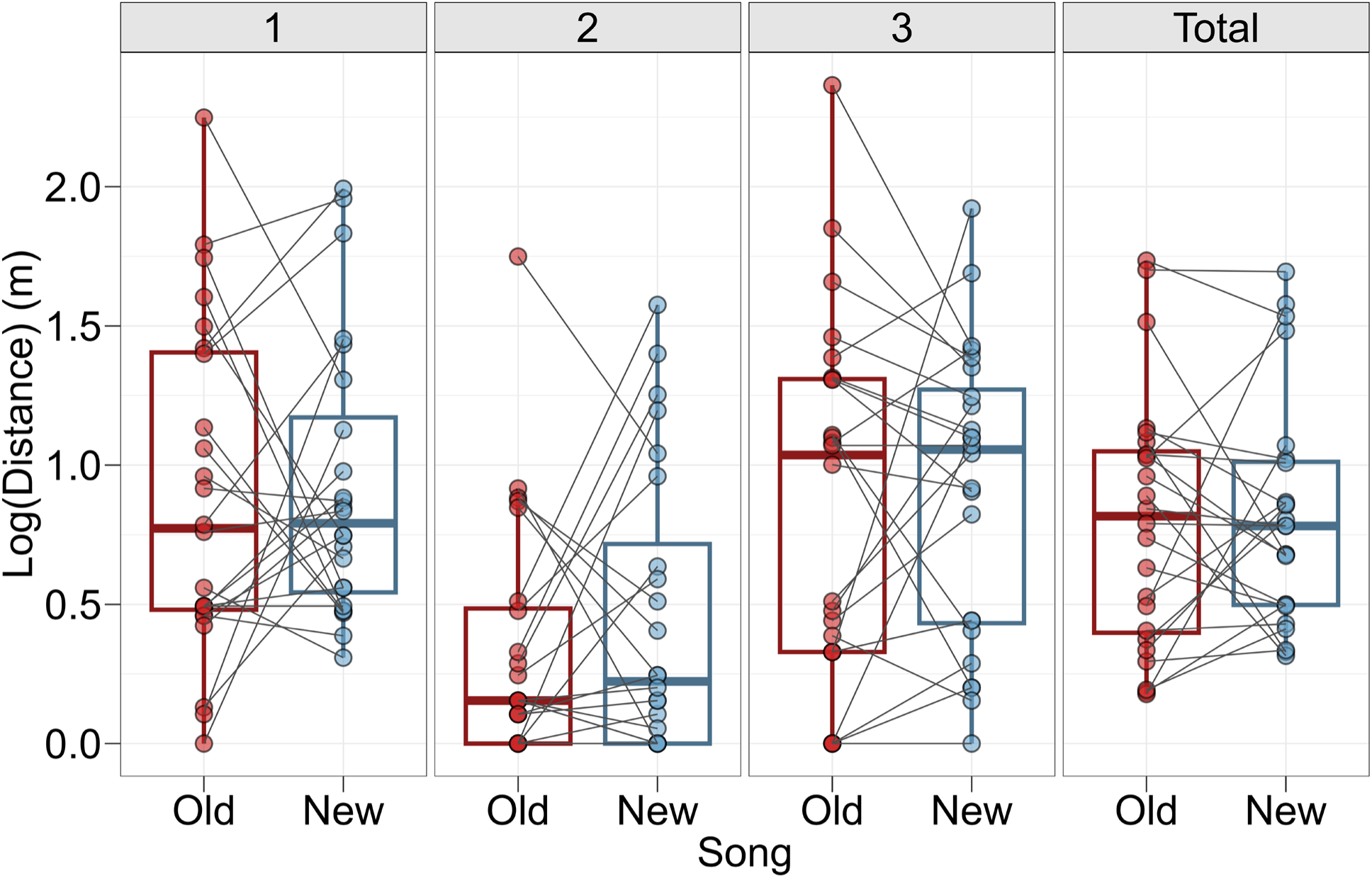
Birds responded similarly to historical (Old) (1992-1994) and current (New) (2021) songs. A closer approach indicates a stronger response, and lines connect responses of the same individuals (within-subjects design). Results are given separately for the first three-minutes of playback (1), the second three-minutes of playback (2), a three-minute post-playback period (3), and for the total trial period (Total). Boxplots are in the style of Tukey: thick lines represent medians, lower and upper extremities of the box are the 25% and 75% quantiles, respectively, and whiskers extend to 1.5 interquartile range past the 25% and 75% quantiles.

**Table 1.**
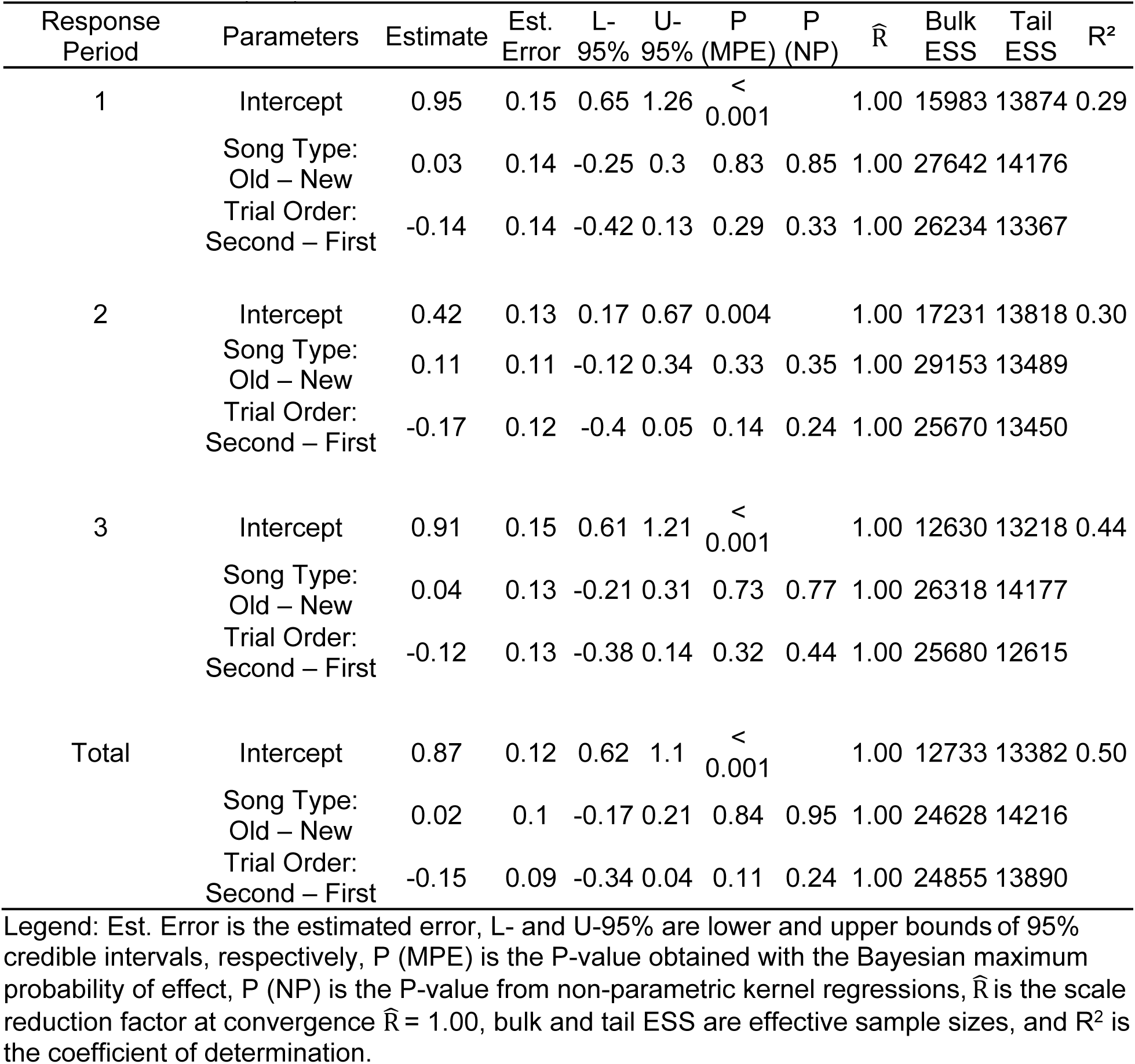
Results of the Bayesian multilevel models and P-values of non-parametric model in the three 3-min periods (1–3) and total trial time.

| Response Period | Parameters | Estimate | Est. Error | L- 95% | U- 95% | P (MPE) | P (NP) | $\hat{R}$ | Bulk ESS | Tail ESS | R<sup>2</sup> |
| --- | --- | --- | --- | --- | --- | --- | --- | --- | --- | --- | --- |
| 1 | Intercept | 0.95 | 0.15 | 0.65 | 1.26 | < 0.001 |  | 1.00 | 15983 | 13874 | 0.29 |
|  | Song Type: Old – New | 0.03 | 0.14 | -0.25 | 0.3 | 0.83 | 0.85 | 1.00 | 27642 | 14176 |  |
|  | Trial Order: Second – First | -0.14 | 0.14 | -0.42 | 0.13 | 0.29 | 0.33 | 1.00 | 26234 | 13367 |  |
| 2 | Intercept | 0.42 | 0.13 | 0.17 | 0.67 | 0.004 |  | 1.00 | 17231 | 13818 | 0.30 |
|  | Song Type: Old – New | 0.11 | 0.11 | -0.12 | 0.34 | 0.33 | 0.35 | 1.00 | 29153 | 13489 |  |
|  | Trial Order: Second – First | -0.17 | 0.12 | -0.4 | 0.05 | 0.14 | 0.24 | 1.00 | 25670 | 13450 |  |
| 3 | Intercept | 0.91 | 0.15 | 0.61 | 1.21 | < 0.001 |  | 1.00 | 12630 | 13218 | 0.44 |
|  | Song Type: Old – New | 0.04 | 0.13 | -0.21 | 0.31 | 0.73 | 0.77 | 1.00 | 26318 | 14177 |  |
|  | Trial Order: Second – First | -0.12 | 0.13 | -0.38 | 0.14 | 0.32 | 0.44 | 1.00 | 25680 | 12615 |  |
| Total | Intercept | 0.87 | 0.12 | 0.62 | 1.1 | < 0.001 |  | 1.00 | 12733 | 13382 | 0.50 |
|  | Song Type: Old – New | 0.02 | 0.1 | -0.17 | 0.21 | 0.84 | 0.95 | 1.00 | 24628 | 14216 |  |
|  | Trial Order: Second – First | -0.15 | 0.09 | -0.34 | 0.04 | 0.11 | 0.24 | 1.00 | 24855 | 13890 |  |

Legend: Est. Error is the estimated error, L- and U-95% are lower and upper bounds of 95% credible intervals, respectively, P (MPE) is the P-value obtained with the Bayesian maximum probability of effect, P (NP) is the P-value from non-parametric kernel regressions, $\hat{R}$ is the scale reduction factor at convergence $\hat{R} = 1.00$ , bulk and tail ESS are effective sample sizes, and R<sup>2</sup> is the coefficient of determination.

## Discussion

We did not find evidence that song sparrows discriminate between songs that had been recorded from the same population approximately 28 years apart. Similar experimental designs, measuring aggressive response in territorial males, have documented discrimination of changes in song over similar time periods in white-crowned sparrows (Derryberry 2007; Derryberry 2011; Luther and Derryberry 2012) and savannah sparrows (Williams et al. 2022) but not in thrush nightingales (Ivanitskii et al. 2025). In both white-crowned and savannah sparrows, females also have been shown to discriminate between current and historical songs (Derryberry 2007; Williams et al. 2022). Given that both song sparrows and thrush nightingales have much greater within-population song diversity than either white-crowned sparrows or savannah sparrows, these results overall support the hypothesis that high within-population song diversity hinders discrimination of cultural evolutionary change in birdsong.

Male song sparrows failed to discriminate historical songs from current songs despite the fact that cultural change had occurred in the song population between our historical and current periods, as shown by our song similarity analysis. The relevant finding was that the introductory phrases of the current playback song types are more common in the current population than are the introductory phrases of the historical playback songs. Thus, it seems possible that a song sparrow could use knowledge of the introductory phrases present in the current song population to judge whether or not a song it hears is typical of the current population. One problem with this proposal is that there appear to be a great many different introductory phrases in a song sparrow population, almost certainly on the order of hundreds (Borror 1961), making the task of learning and remembering a sufficient sample of the introductory phrases very difficult. Another problem is that it is not clear that the familiarity of component phrases affects the response of song sparrows to song; Searcy et al. (2003) present a counter-example, where substituting a highly familiar phrase into foreign songs decreased, rather than increased, the response of male song sparrows to those songs. Either or both of these problems could explain why song sparrows failed to discriminate between current and historical songs in our playback experiment.

Discrimination of current from historical song may be adaptive in some cases of song evolution, especially ones in which songs have evolved adaptively. In an urban population of white-crowned sparrows, for example, songs evolved towards higher minimum frequencies as low frequency traffic noise became more intense (Luther & Derryberry 2012). Under these conditions, high frequency signals suffer less masking (Luther & Derryberry 2012) and increase in relative abundance perhaps in part because unmasked and undegraded songs are preferentially learned by young males (Peters et al. 2012, Moseley et al 2018). Similarly, selection on trill characteristics due to changes in vegetation density may explain rapid song evolution in other white-crowned sparrow populations (Derryberry 2009) and hence explain at least in part greater receiver response to current than historical songs (Derryberry 2007). In both these white-crowned sparrow cases, stronger response to current songs seems to be explained by those songs being better adapted to transmission under current environmental conditions. In the savannah sparrow studies, certain song elements spread through the population because songs with those elements were preferentially copied by young males (Williams et al. 2022). Young males may prefer to learn songs with these elements because such songs are preferred by females (Williams et al. 2022), leading to higher reproductive success in males that sing them (Williams et al 2013). Here, males singing current songs may be more serious rivals, making heightened aggressive response to current songs adaptive.

Cultural evolution of song is often suggested to occur through the cumulative effect of errors in song learning (Slater 1986; Podos and Warren 2007; Catchpole and Slater 2008), in which case song evolution would in general not be in the direction of greater adaptation. Thus, it is not surprising that in most known cases of short-term song evolution, there is no obvious adaptive value to the observed changes in song. The absence of known adaptive effects particularly applies to the many cases in which song types or syllable types increase or decrease in relative abundance over one or a few decades (Payne 1985; Nelson et al. 2004; Byers et al. 2010; Ju et al. 2019; Souriau et al. 2025). Also suspect are cases in which average acoustic measures change over time in a way that is not known to track environmental changes (Derryberry 2011; Garcia et al. 2015). In cases in which temporal changes in song have no adaptive consequences for signalers, it is difficult to see why discriminating current from historical songs would be adaptive for receivers. The temporal changes we document in our song sparrow population in the occurrence of some introductory phrases seem likely to fall into the category of non-adaptive changes, suggesting that the failure of our subjects to respond to these changes is not maladaptive.

The same processes of cultural evolution that produce changes in song over time in a single location presumably are also responsible for producing differences in song over space at a single time (Podos and Warren 2007; Catchpole and Slater 2008). Just as we would expect differences between temporal samples to increase with the length of time between them, so we would expect differences between geographic samples to increase with the spatial distance between them. This reasoning suggests that there may an approximate relationship between temporal and geographic differences, such that a change in songs at one location over a certain specified number of years is approximately equivalent to a difference in contemporary songs over a certain specified geographic distance. Derryberry (2011) examined this kind of relationship in white-crowned sparrows, using male response to territorial playback to gauge changes in song over time and space. Her results indicate that changes in white-crowned sparrow song within a locality over 27 years are approximately equivalent to differences between adjacent dialects within a single time and are considerably less than the differences between non-adjacent dialects. In our song sparrow work, we have found that males discriminate between local western Pennsylvania songs and songs from New York sites approximately 540 km distant (Searcy et al. 1997) but not between local songs and foreign songs from half that distance (∼270km) or less (135, 68, 34, and18 km) (Searcy et al. 2002). Thus, at this point we can only say that the temporal differences arising in ∼28 years seem to be less than the geographic differences existing over a distance of ∼540 km.

## Acknowledgements

We thank Cori Richards-Zawacki and the staff of the Pymatuning Laboratory of Ecology, University of Pittsburgh, for logistical support, and the Pennsylvania Game Commission and Pymatuning State Park for granting access to study sites. We also thank Jill Soha (Department of Biology, Duke University) for assistance with preparing playback stimuli. Funding was provided by Duke University’s Trinity College of Arts and Sciences.

## Data availability

The experimental data that support the findings of this study are available in Figshare with the identifier https://doi.org/10.6084/m9.figshare.32863079

## Declarations

### Ethical approval

All procedures were approved by the Institutional Animal Care and Use Committees of Duke University (protocol #A182-22-10) and the University of Pittsburgh (protocol #24044976). All applicable national and institutional guidelines for the ethical treatment of animals were followed. The research was conducted with free-living birds that were not captured or otherwise handled for this study. Any disturbance associated with conducting a playback trial lasts less than one-half hour, including time needed to set up and remove the equipment and distance markers used to conduct the trial. Individuals were only tested twice, with tests separated by 2 to 4 days

